# Characterization of the *Rhizophagus irregularis* multicopper oxidase family indicates that the iron transporter RiFTR1 does not require a ferroxidase partner

**DOI:** 10.1101/2022.06.14.496117

**Authors:** E Tamayo, C Shim, AG Castillo, JP Benz, N Ferrol

## Abstract

The contribution of arbuscular mycorrhizal fungi (AM fungi) to plant iron (Fe) acquisition has been demonstrated in several studies. Recently, it has been shown that AM fungi use a high-affinity reductive pathway for Fe uptake. In the AM fungus *Rhizophagus irregularis* the ferric reductase RiFRE1 and the Fe permeases RiFTR1 and RiFTR2 have already been characterized. In an attempt to identify the third component of the reductive iron uptake pathway, a genome-wide approach has been used in *R. irregularis* to find genes encoding ferroxidases of the multicopper oxidase (MCO) gene family. Nine genes putatively encoding MCOs (*RiMCO1-9*) were identified. A phylogenetic analysis of MCO sequences of fungi from different taxonomic groups revealed that all RiMCOs clustered together in the ferroxidase/laccase group, and none with the Fet3-type ferroxidases. *RiMCO1* and *RiMCO3* were the only *MCO* genes displaying a detectable gene expression pattern typical of a high-affinity Fe transport system, indicating that RiMCO1 and RiMCO3 might have a role in the reductive high-affinity Fe uptake system. Moreover, yeast mutant complementation assays showed that the iron permease RiFTR1 can operate without the presence of a ferroxidase, indicating that it is able to transport also ferrous (II) iron.

## Introduction

Iron (Fe) is an essential metal element, however, it can become extremely toxic at high concentrations. Therefore, its biological levels are finely regulated in living cells. To maintain Fe homeostasis, all organisms have evolved sophisticated tools to cope with their requirements for it. Although Fe is abundant in nature, it has a low availability because it is most commonly found as ferric (III) hydroxide, which is a rather stable and poorly soluble compound (Kosman, 2003). For this reason, high-affinity Fe transport systems are required for uptake.

The symbioses with arbuscular mycorrhizal (AM) fungi belonging to the Glomeromycotina (Mucoromycota phylum) is an evolutionary ancient strategy of plants to increase their nutrient supply. AM fungi are obligate biotrophs that form extensive external mycelium in the soil that can absorb nutrients beyond the depletion zone that develops around the roots. This way, they provide a new pathway for the uptake and transport of low mobility nutrients in the soil, particularly P and N, but also micronutrients such as Fe, Cu and Zn (Smith and Read, 2008). In return, the plants provide carbohydrates and lipids to the fungi (Shachar-Hill *et al*., 1995; Helber *et al*., 2011; Keymer *et al*., 2017; Luginbuehl *et al*., 2017).

The contribution of AM fungi to plant Fe uptake has been shown in several studies (for example, Caris *et al*., 1998; Kobae *et al*., 2014; Kabir *et al*., 2020) and we recently demonstrated that AM fungi use a high-affinity Fe reductive pathway to take up Fe from the soil (Tamayo *et al*., 2018). As mentioned above, Fe is mostly found in insoluble Fe(III) complexes in soils. In the well characterized yeast *Saccharomyces cerevisiae*, the high-affinity reductive Fe uptake system thus requires reduction of Fe(III) to Fe(II) by membrane-bound ferrireductases, which is then rapidly internalized by the concerted action of a ferroxidase (FET3) and an iron permease (FTR1) that form a plasma membrane protein complex (Askwith *et al*., 1994; Stearman *et al*., 1996; Kosman, 2010). Fe(II) is first oxidized by FET3, and then transported into the cytosol as Fe(III) by FTR1 via a channeling mechanism (Kwok *et al*., 2006). In our previous work, we identified two components of this high-affinity iron uptake system in the AM fungus *R. irregularis*, the ferric reductase RiFRE1 and the Fe permeases RiFTR1 and RiFTR2 (Tamayo *et al*., 2018). However, the ferroxidase partners of the two iron permeases remained to be identified.

Fungal ferroxidases (Fe(II): oxygen oxidoreductase, EC 1.16.3.1) are plasma membrane proteins with a single transmembrane domain and an extracellular multicopper oxidase domain responsible for the ferroxidase activity. They belong to the family of multicopper oxidases (MCOs), proteins that have the ability to couple the oxidation of a variety of substrates with the reduction of molecular oxygen to water using four copper atoms (Rydén and Hunt, 1993). Other members of this family are laccases, ascorbate oxidases, bilirubin oxidases and potential ferroxidases/laccases exhibiting either one or both enzymatic functions (Hoegger *et al*., 2006; Gräff *et al*., 2020). Fungal Fet3-like ferroxidases have been characterized in many fungi including *Schizosaccharomyces pombe, Pichia pastoris, Arxula adeninivorans, Aspergillus fumigatus, Ustilago maydis, Fusarium graminearum, Phanerochaete chrysosporium,Cryptococcus neoformans, Candida albicans* and *Colletotrichum graminicola* (Askwith and Kaplan, 1997; Ahmad *et al*., 2022; Paronetto *et al*., 2001; Wartmann *et al*., 2002; Schrettl *et al*., 2004; Eicchorn *et al*., 2006; Greenshields *et al*., 2007; Larrondo *et al*., 2007; Jung *et al*., 2009; Ziegler *et al*., 2011; Albarouki and Deising, 2013), while the ferroxidases/laccases have been characterized in the fungi *P. chrysosporium* (Larrondo *et al*. 2003) and *C. neoformans* (Liu *et al*., 1999).

With the aim of identifying the ferroxidase partners of the iron permeases of *R. irregularis*, a genome-wide approach has been used to find candidate genes belonging to the MCO gene family. Nine genes putatively encoding MCOs were identified. Phylogenetic analyses revealed that all RiMCOs grouped in the ferroxidase/laccase cluster of MCOs. *In silico* and expression analyses indicate that RiMCO1 and RiMCO3 have the potential to represent the ferroxidase partners of the two Fe permeases of the reductive pathway, but heterologous expression and growth assays suggest that the Fe transporter RiFTR1 may act without the assistance of a ferroxidase.

## Experimental procedures

### *Identification of* MCO *genes in the* R. irregularis *genome and sequence analyses*

Amino acid sequences of fungal MCOs were retrieved from the freely accessible transport database TCDB (http://www.tcdb.org/). These sequences were used to search for orthologous sequences in the filtered model dataset of *R. irregularis* on the JGI website (http://genome.jgi-psf.org/Gloin1; Tisserant *et al*., 2013) using Basic Local Alignment Search Tool (BLAST) algorithm (Altschul *et al*., 1990) via a protein BLAST. A second search was performed via a keyword search directly using “multicopper oxidase”, “laccase”, “ascorbate oxidase”, “ferroxidase”, “bilirubin oxidase” and “Fet3” as keywords. Since some of the fungal reference proteins were phylogenetically distant from *R. irregularis*, manually curated *Laccaria bicolor* (http://genome.jgi-psf.org/Lacbi2), *Tuber melanosporum* (http://genome.jgi.doe.gov/Tubme1) and *Rhizopus oryzae* (http://genome.jgi.doe.gov/Rhior3) databases were used to look for additional orthologous sequences in the filtered model dataset of *R. irregularis*. This was also done via a BLASTp, run with the standard program settings. Finally, a keyword search for putative multicopper oxidases was performed on the functional annotation collection from Lin *et al*. (2014). Sequences were finally compared to the most recent version delivered sequences of *R. irregularis* via BLASTp and keyword search for putative multicopper oxidases (https://genome.jgi.doe.gov/RhiirA1_1/RhiirA1_1.home.html; https://genome.jgi.doe.gov/RhiirB3/RhiirB3.home.html; https://genome.jgi.doe.gov/RhiirC2/RhiirC2.home.html; https://genome.jgi.doe.gov/RhiirA4/RhiirA4.home.html; https://genome.jgi.doe.gov/RhiirA5/RhiirA5.home.html; https://genome.jgi.doe.gov/Rhiir2_1/Rhiir2_1.home.html; Chen *et al*., 2018).

Predictions of putative transmembrane domains were made using the TMHMM Server v.2.0 (http://www.cbs.dtu.dk/services/TMHMM/) and the presence of the conserved domains was analyzed using SMART software (http://smart.embl-heidelberg.de/) (Letunic *et al*. 2021).

Full-length amino acid sequences were aligned with the orthologous sequences of a number of fungi representatives of distinct taxonomic groups by ClustalW (Version 2.1; Larkin *et al*., 2007; http://www.ebi.ac.uk/Tools/msa/clustalw2/) with default setting. Alignments were imported into the Molecular Evolutionary Genetics Analysis (MEGA) package version 11 (Tamura *et al*., 2021). Phylogenetic analyses were conducted by the Neighbor-Joining (NJ) method, implemented in MEGA, with a pair-wise deletion of gaps and the Poisson model for distance calculation. Bootstrap analyses were carried out with 1000 replicates. The evolutionary tree was drawn to scale.

### Biological materials and growth conditions

*Rhizophagus irregularis* monoxenic cultures were established as described by St-Arnaud *et al*. (1996), with some modifications. Briefly, clone DC2 of carrot (*Daucus carota* L.) Ri-T DNA transformed roots were cultured with the AM fungus *R. irregularis* Schenck and Smith DAOM 197198 in two-compartment Petri dishes. Cultures were initiated in one compartment of each plate containing M medium (Chabot *et al*., 1992) by placing several non-mycorrhizal carrot root segments and a piece of fungal inoculum containing extraradical mycelium (ERM), fragments of mycorrhizal roots and spores. Fungal hyphae and roots were allowed to grow over to the other compartment containing the same M medium. Plates were incubated in the dark at 24 °C for 7–8 weeks until the second compartment was profusely colonized by the fungus and the roots. Then, the older compartment was removed and refilled with liquid M medium without sucrose (M-C medium) containing different Fe concentrations: 0.045 mM (control), 4.5 mM or 45 mM EDTA Fe(III) sodium salt. Fungal hyphae, but not roots, were allowed to grow over to this compartment (hyphal compartment). Plates were incubated in the dark at 24 °C for 2–3 additional weeks. To induce Fe-deficient conditions, monoxenic cultures were initiated in plates containing solid M medium without Fe and incubated in the dark for 7-8 weeks as above. Later, fungal hyphae grown (for 2-3 weeks) in liquid M-C medium without Fe were exposed 3 days to 0.5 mM ferrozine (Sigma). This was done by replacing the liquid medium by 20 ml of a freshly prepared liquid M-C medium without Fe and supplemented with 0.5 mM ferrozine, which was prepared by adding 0.2 ml of a 0.5 mM stock solution in 50 mM MES pH 6.1 (ferrozine M media). Control plates were prepared as described above, but the M-C media used to refill the hyphal compartment was supplemented with 2 ml 50 mM MES pH 6.1. Plates were incubated in the dark at 24 °C for 3 additional days.

ERM from the different hyphal compartments was directly recovered under sterile conditions by using a pair of tweezers, washed with sterile water and dried on filter paper. The mycelium was immediately frozen in liquid nitrogen and stored at -80 °C until used.

To analyze intraradical gene expression, hyphae growing in the hyphal compartment were used as a source of mycorrhizal inoculum. Carrot roots were placed on top of a densely colonized hyphal compartment and collected 15 days later. Extraradical hyphae attached to the roots were removed with forceps under a binocular microscope. Roots were then frozen in liquid N and stored at -80°C until used.

The *S. cerevisiae* strains used in this study are listed in Table S1. Strains were grown on YPD or complete synthetic medium (CSM), supplemented with appropriate amino acids.

### Nucleic acid extraction and cDNA synthesis

Total fungal RNA from ERM from the different treatments of *R. irregularis* and mycorrhizal carrot roots was extracted using the RNeasy Plant Mini Kit (QIAGEN, Maryland), following manufacturer’s instructions. DNase treatment was performed using RNA-free DNase Set (QIAGEN, Maryland) following the manufacturer’s instructions. cDNAs were obtained from 1 µg of total DNase-treated RNA in a 20 µl reaction containing 200 units of Super-Script III Reverse Transcriptase (Invitrogen) and 50 pmol oligo (dT)_20_ (Invitrogen), according to the manufacturer’s instructions.

### Gene isolation

The 5’ end of *RiMCO1* was verified by rapid amplification of cDNA ends (RACE) using the SMART RACE cDNA amplification kit (Clontech, Palo Alto, CA, USA), the *RiMCO1*-specific primer GiFETa.rR (Table S2) and 1 µg total RNA from ERM grown in control plates. The full-length cDNA of *RiMCO1* was obtained by PCR amplification of *R. irregularis* cDNA, using the corresponding primer pairs: RiFETa_s.fF and FETa.fR (Table S2). The full-length sequence of *RiMCO3* was obtained by RACE using the *RiMCO3*-specific primers FETc.rF and FETc.rR for the 3’and 5’RACE reactions, respectively, designed based on the sequence GenBank Accession No. AUPC01010373, and 1 µg total RNA from ERM grown in control plates. Full-length cDNA of *RiMCO3* was obtained by PCR amplification of *R. irregularis* cDNA, using the corresponding primer pairs: FETc.fF and RiFETc_l.fR (Table S2). PCR products were cloned in the pGEM-T easy vector (Promega, Madison, USA).

All plasmids were amplified by transformation of *E. coli* following standard procedures and purified by using the Qiagen Miniprep Kit (Qiagen, Maryland, USA). All sequences and constructs were checked by sequencing before further use. Nucleotide sequences were determined by Taq polymerase cycle sequencing by using an automated DNA sequencer (ABI Prism 3130xl Genetic Analyzer, Applied Biosystems, Carlsbad, USA).

### Heterologous expression

For heterologous gene expression analyses, the *RiFTR1, RiFTR2, RiMCO1* and *RiMCO3* full-length cDNAs were obtained by PCR amplification of the genes cloned in pGEM-T easy vector (Promega, Madison, USA), using the corresponding primer pairs (Table S1). The RIFTR1::pGEM-T and RiFTR2:: pGEM-T plasmids were obtained in a previous work (Tamayo *et al*., 2018). PCR products were cloned in the part entry vector pYTK001 via BsmBI Golden Gate assembly (Lee *et al*., 2015). All plasmids were amplified by transformation of *E. coli* following standard procedures and purified by using the Hi Yield® Plasmid Mini DNA Isolation Kit (SLG®). All sequences and constructs were checked by sequencing before further use. For cassette assemblies, the cassette entry vector pYTK095 was used. *RiFTR1* and *RiFTR2* were cloned via BsaI-v2 assembly with the promoter of *CCW12* and the terminator of *PGK1*, and *RiMCO1* and *RiMCO3* with the promoter of *TDH3* and the terminator of *ADH1*. Multicassettes were constructed via BsmBI assembly using the integration vector pYTK096 and single cassettes or the combination of a pair of FTR-MCO cassettes. The full-length cDNAs of *ScFtr1* and *ScFet3* were also amplified and multicassettes were constructed and used as positive controls in the complementation analyses. In total, nine multicassettes were built: *RiFTR1, RiFTR2, RiFTR1-RiMCO1, RiFTR2-RiMCO1, RiFTR1-RiMCO3, RiFTR2-RiMCO3, ScFtr1, ScFet3* and *ScFtr1-ScFet3* muticassettes.

Yeasts were transformed with the corresponding constructs using a lithium acetate-based method (Schiestl and Gietz, 1989), and transformants (Table S1) were selected in CSM medium by auxotrophy to uracil.

For the complementation assay, a specially designed medium was used (Knight *et al*., 2002). To avoid Fe contamination, new plastic or acid-washed glassware was used for all media preparation. Briefly, the assay medium consisted of CSM made with a yeast nitrogen base lacking Fe and copper (Bio-101), supplemented with 1 mM copper sulfate and 50 mM MES pH 6.1. The Fe-limited medium was the assay medium with 100 µM bathophenanthrolinedisulfonate (BPS) (Sigma) (10 mM stock) and different concentrations of ferrous ammonium sulfate. For the plating assay, the medium contained 2% Bacto agar. Stationary yeast cultures with Δ*fet3* and Δ*ftr1* background grown in CSM liquid culture were washed with ice-cold distilled water, resuspended in Fe-limited medium supplemented with 200 µM ferrous ammonium sulfate to an O.D.=0.1 and grown to saturation overnight. Stationary yeast cultures with Δ*fet3*Δ*fet4* and Δ*fet3*Δ*fet4*Δ*ftr1* background were grown in CSM liquid culture supplemented with 100 µM ferrous ammonium sulfate to saturation overnight. These cultures were then washed twice and diluted to an optical density at 600 nm of 1. Serial 1:5 dilutions were spotted (5 ml) onto plates containing the assay media supplemented with BPS and different Fe concentrations to determine the lowest Fe concentration at which each mutant strain was able to grow.

### Protein localization analyses

Localization of *R. irregularis* FTR1 protein in *S. cerevisiae* was analyzed by fusion of the mRuby2 protein gene to the C-terminus using an *mRuby2*-Part3b plasmid via Golden Gate assembly (Lee *et al*., 2015). *mRuby2* and *ScFtr1-mRuby* were used as negative and positive controls, respectively. The Δ*fet3*Δ*fet4* yeast mutant strain was transformed with the resulting multicassettes. Stationary yeast cultures grown in CSM were washed twice with ice-cold distilled water, diluted in Fe-limited medium supplemented with 200 µM ferrous ammonium sulfate and grown for several hours in this medium to an OD_600_ between 0.8 and 1.6. Cells were washed three times and resuspended in water before visualization. The fluorescence signal was visualized with a Zeiss Axio fluorescent microscope. mRuby2 fusion proteins were imaged using a 542-582 nm filter. Image sets were processed and overlapped using ImageJ (http://fiji.sc/Fiji).

### Gene expression analyses

*RiMCOs* gene expression was studied by real-time RT-PCR by using an iQTM5 Multicolor Real-Time PCR Detection System (Bio-Rad). Each 20 µl reaction contained 1 µl of a 1:10 dilution of the cDNA, 200 nM each primer, 10 µl of iQTM SYBR Green Supermix 2x (Bio-Rad). The PCR program consisted in a 3-min incubation at 95 °C to activate the hot-start recombinant Taq DNA polymerase, followed by 36 cycles of 30 s at 95 °C, 30 s at 58 °C and 30 s at 72 °C, where the fluorescence signal was measured. The specificity of the PCR amplification procedure was checked with a heat-dissociation protocol (from 58 to 95 °C) after the final cycle of the PCR. Primer specificity was checked by sequencing the amplicons obtained with each primer pair. The primer sets used are listed in Table S2. The efficiency of the primer sets was evaluated by performing real-time PCR on several dilutions of cDNA. Because RNA extracted from mycorrhizal roots contains plant and fungal RNAs, specificity of the primer pairs was also analyzed by PCR amplification of genomic DNA isolated from non-mycorrhizal carrot roots and of cDNA from non-colonized carrot roots. The results obtained for the different treatments were standardized to the elongation factor 1-alpha gene levels (GenBank Accession No. DQ282611), which were amplified with the following primers: GintEFfw and GintEFrev. RT-PCR determinations were performed on three independent biological samples from three replicate experiments. Real-time PCR experiments were carried out three times for each biological sample, with the threshold cycle (Ct) determined in triplicate. The relative levels of transcription were calculated by using the 2-^ΔΔ^CT method (Schmittgen and Livak, 2008), and the standard error was computed from the average of the ΔCT values for each biological sample.

### Statistical analyses

Statgraphics Centurion XVI software was used for the statistical analysis of the means and standard deviation determinations. ANOVA, followed by a Fisher’s LSD test (*p*<0.05) when possible, was used for the comparison of the treatments based on at least 3 biological replicates for each treatment (n ≥ 3).

## Results

### R. irregularis *has at least nine multicopper oxidases in its genome*

A search for putative MCOs in the *R. irregularis* genome databases (*R. irregularis* DAOM 181602, A1, B3, C2, A4, A5 and DAOM 197198) led to the identification of nine genes putatively encoding MCOs. Since the length of the RiMCO1, RiMCO3, RiMCO8 and RiMCO9 protein sequences found in the *R. irregularis* genome databases was shorter than expected, we attempted to obtain the full-length cDNAs by RACE. The open reading frames and number of introns of *RiMCO4, RiMCO5* and *RiMCO6* were also confirmed by cloning and sequencing the respective cDNA sequences. However, the coding sequences of *RiMCO2, RiMCO7, RiMCO8* and *RiMCO9* could not be experimentally confirmed because they were expressed below detection levels in the *R. irregularis* ERM. The length of the nucleotide coding sequences of the nine *RiMCO* genes ranged from 1620 to 2127 bp, while the length of the nucleotide genomic sequences varied from 2159 to 2836 bp (Table S3). Comparison between the cDNA and genomic sequences revealed that the number of introns in the individual *RiMCOs* varies from 4 to 12 (Table S3 and Fig. S1).

The length of the predicted RiMCO proteins ranges from 517 to 708 amino acids (Table S3) and they contain three cupredoxin-like domains linked by external interdomains (Fig. 1A). Sequence identities among the deduced amino acid sequences of RiMCO1-9 ranges from 24 to 82%, with RiMCO6 having the lowest identity to the others (≤29%) (Fig. S2). RiMCO4 and RiMCO7 showed the highest degree of identity (82%). Relationships between the nine *R. irregularis* MCOs are also reflected in the phylogenetic tree based on their amino acid similarities (Fig. 1B). Based on the similarity of the RiMCO gene products and on the intron positions, five gene subfamilies were defined (Figs. 1B and S1).

**Figure 1.**
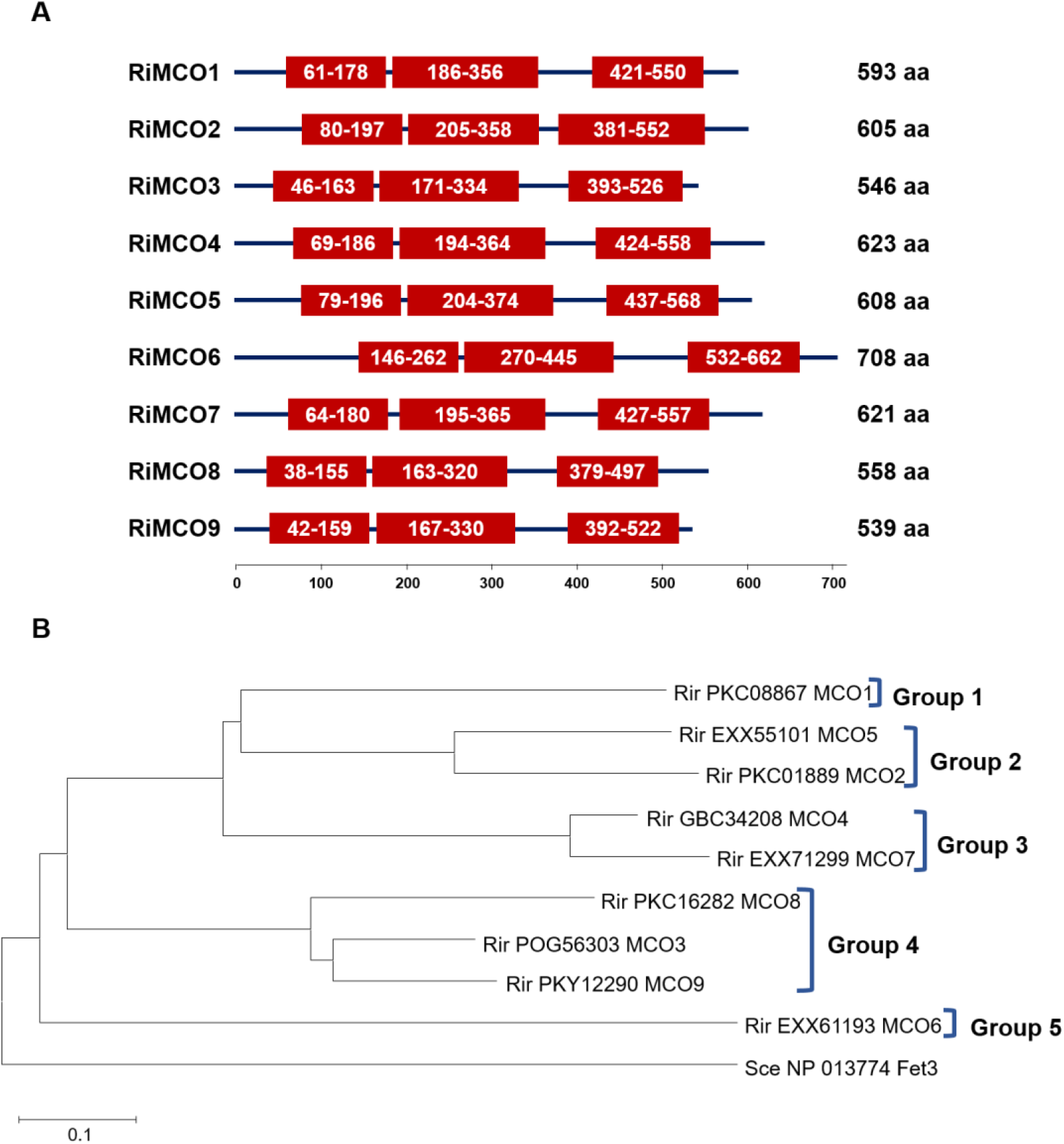
**A. Domain organization in the *Rhizophagus irregularis MCO* proteins.** Domains were identified by SMART analyses. Red boxes indicate the cupredoxin-like domains. Numbers represent amino acid positions. Protein lengths in amino acids (aa) are indicated on the right. **B. Neighbor-Joining tree of the deduced amino acid sequences of the *R. irregularis* MCOs**. The *S. cerevisiae* Fet3 sequence was used as outgroup. Rir, *R. irregularis*; Sce, *S. cerevisiae*. Protein NCBI identification numbers are indicated.

Alignment of the deduced amino acid sequences of RiMCO1-9 with other fungal MCOs revealed that the nine RiMCOs have the four conserved regions that contain the residues involved in copper coordination, a characteristic signature of all MCOs (Fig. 2). However, they lack the cysteine in signature sequence S2, which is always present in laccases *sensu stricto* but not in Fet3-type ferroxidases (Kumar *et al*., 2003).

**Figure 2.**
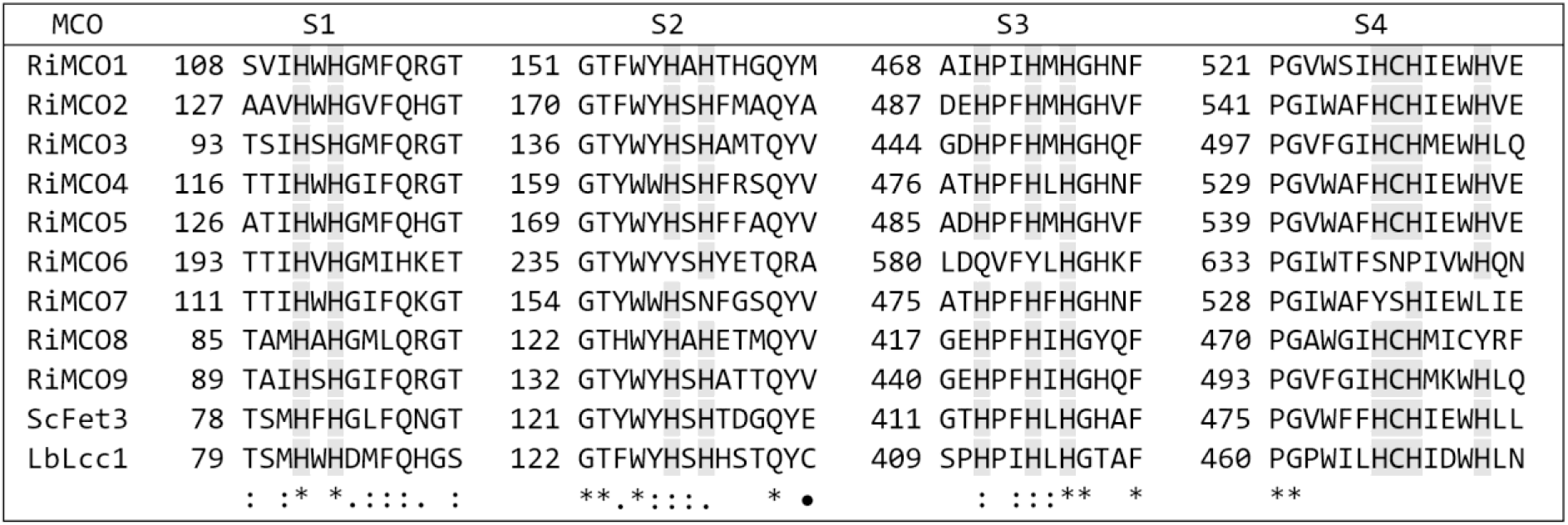
Sequence alignment of the four copper-binding sites (signature motifs S1 to S4) of the nine *Rhizophagus irregularis* MCOs, the *Saccharomyces cerevisiae* ferroxidase Fet3 and the *Laccaria bicolor* laccase Lcc1. Histidine (H) and cysteine (C) copper ligands are indicated. [The conserved 10 histidines and the cysteine residue involved in the coordination of copper are shown]. A black circle denotes a residue in signature sequence S2 that is always a cysteine (C) in classical laccases, but not in Fet3 ferroxidases.

A phylogenetic analysis of MCO sequences of different taxonomic groups revealed that all RiMCOs clustered together in the ferroxidase/laccase group (Fig. 3A and B). In contrast to the MCOs of most other fungi, none of the RiMCOs clustered with the Fet3-type ferroxidases (Fig. 3B). The same is true also for the arbuscular mycorrhizal fungi *Rhizophagus cerebriforme* and *Rhizophagus clarus*, both belonging to the Glomeromycotina subphylum and whose genomes have recently been sequenced (Morin *et al*., 2019).

**Figure 3.**
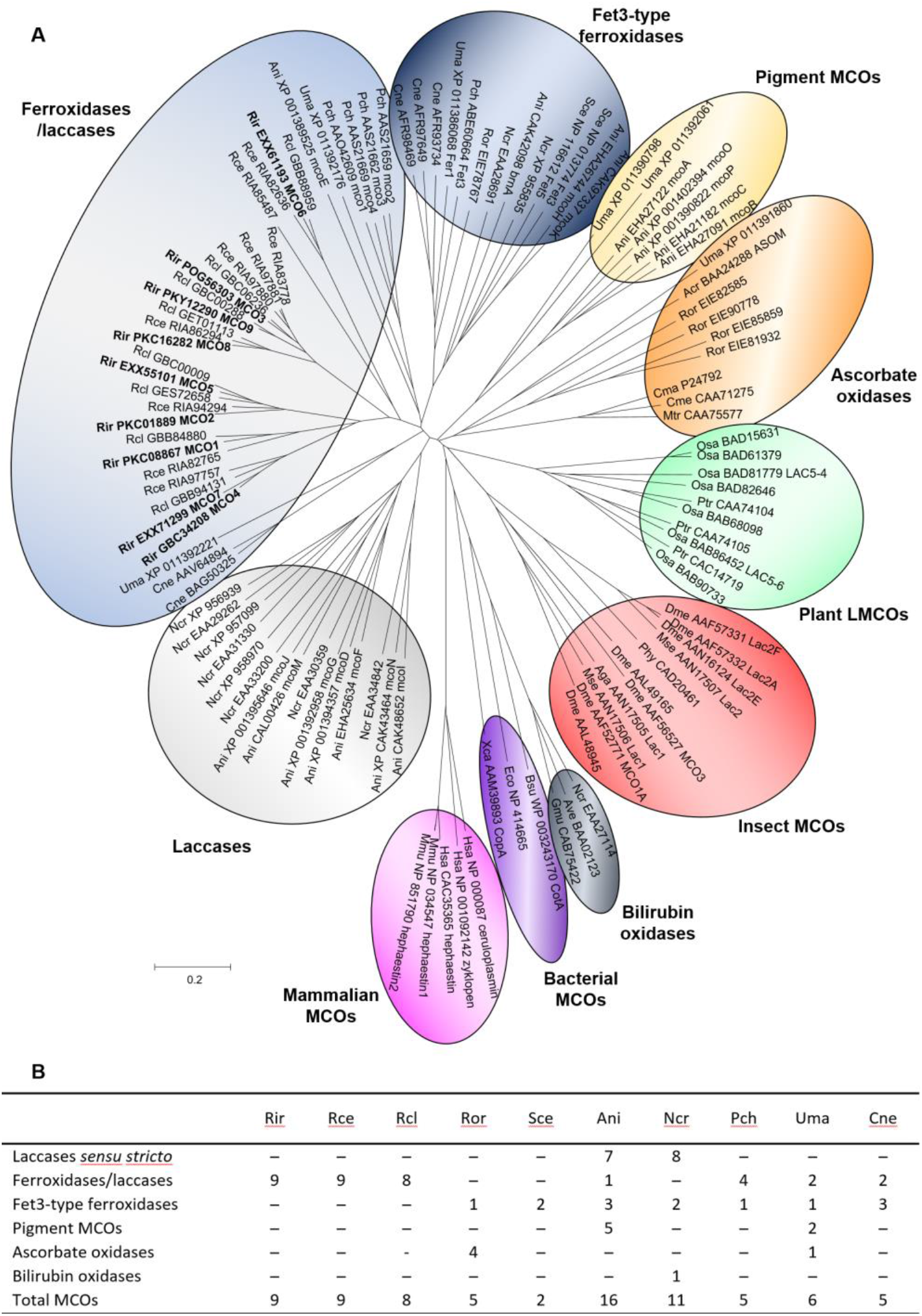
**A. Unrooted Neighbor-Joining tree of the MCO gene family.** Organisms: Acr, *Acremonium* sp.; Aga, *Anopheles gambiae*; Ani, *Aspergillus niger*; Ave, *Albifimbria verrucaria*; Bsu, *Bacillus subtilis*; Cma, *Cucurbita maxima*; Cme, *Cucumis melo*; Cne, *Cryptococcus neoformans*; Dme, *Drosophila melanogaster*; Eco, *Escherichia coli*; Gmu, *Gliomastix murorum*; Hsa, *Homo sapiens*; Mmu, *Mus musculus*; Mse, *Manduca sexta*; Mtr, *Medicago truncatula*; Ncr, *Neurospora crassa*; Osa, *Oryza sativa*; Pch, *Phanerochaete chrysosporium*; Phy, *Pimpla hypochondriaca*; Ptr, *Populus trichocarpa*; Rce, *Rhizophagus cerebriforme*; Rcl, *Rhizophagus clarus*; Rir, *Rhizophagus irregularis*; Ror, *Rhizopus oryzae*; Sce, *Saccharomyces cerevisiae*; Uma, *Ustilgo maydis*; Xca, *Xanthomonas campestris. Rhizophagus irregularis* MCOs are emphasized in bold. Protein NCBI identification numbers are indicated. **B. Number and classification of MCOs in fungal genomes**. Organisms: Ani, *Aspergillus niger*; Cne, *Cryptococcus neoformans*; Ncr, *Neurospora crassa*; Pch, *Phanerochaete chrysosporium*; Rce, *Rhizophagus cerebriforme*; Rcl, *Rhizophagus clarus*; Rir, *Rhizophagus irregularis*; Ror, *Rhizopus oryzae*; Sce, *Saccharomyces cerevisiae*; Uma, *Ustilgo maydis*.

RiMCO1, RiMCO3, RiMCO6, RiMCO8 and RiMCO9 have at least one of the four residues (E185, D283, Y354 and D409) that were shown in *S. cerevisiae* to be essential for Fe oxidation (Wang *et al*., 2003; Stoj *et al*., 2006; Fig. 4), a characteristic of ferroxidases and ferroxidases/laccases with a main ferroxidase activity (Kües and Rühl, 2011).

**Figure 4.**
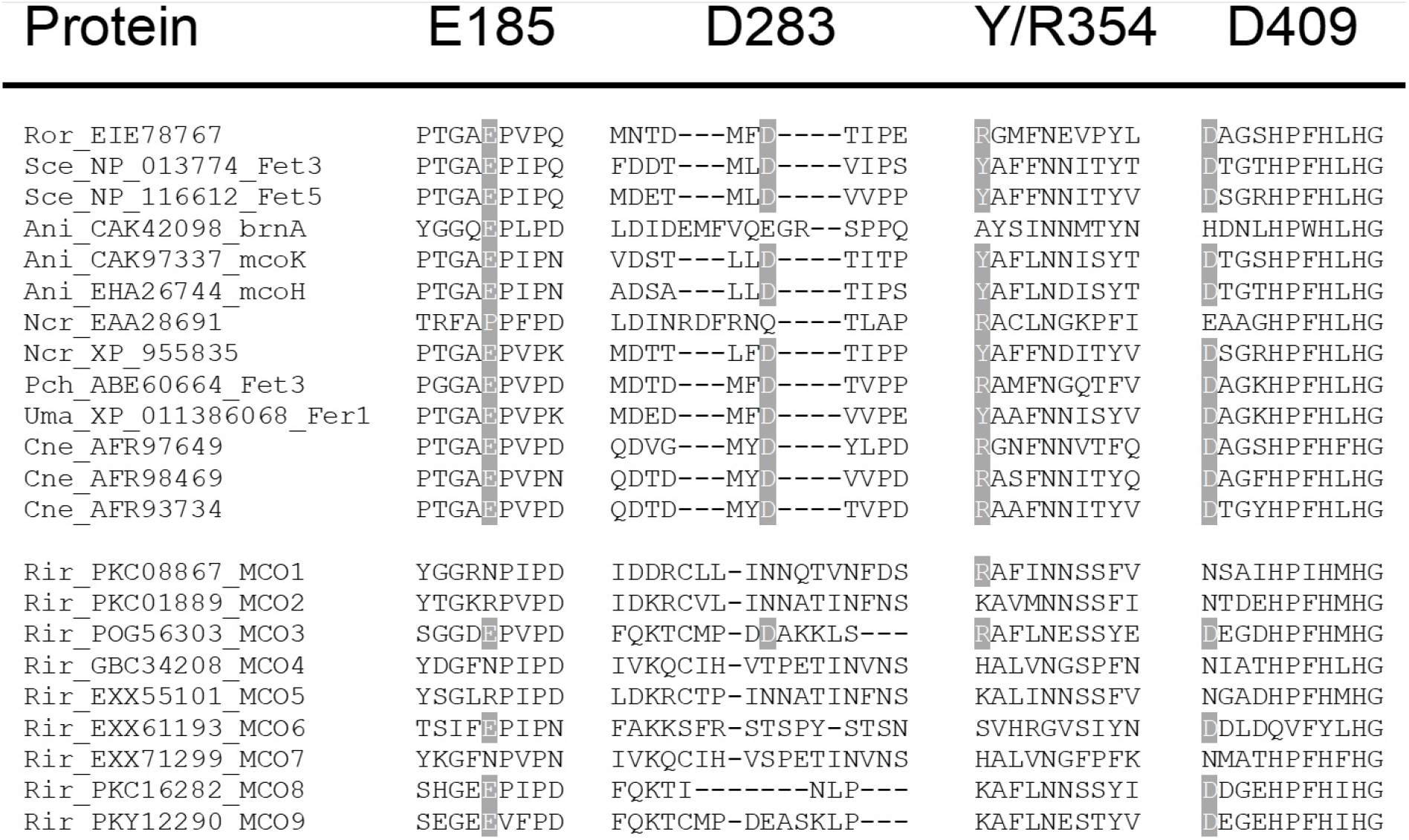
Alignment of the four regions known to function in oxidation of Fe(II) to Fe(III) in *S. cerevisiae* Fet3 with the corresponding sequence regions of Fet3-type ferroxidases from fungi and *Rhizophagus irregularis* MCOs. The upper block presents the cluster of Fet3-like enzymes following the order of the figure 3B and the lower block the cluster of *R. irregularis* ferroxidases/laccases. The residues involved in ferroxidase activity (E185, D283, Y/R354 and D409) are highlighted in grey. Organisms: Ani, *Aspergillus niger*; Cne, *Cryptococcus neoformans*; Ncr, *Neurospora crassa*; Pch, *Phanerochaete chrysosporium*; Rir, *Rhizophagus irregularis*; Ror, *Rhizopus oryzae*; Sce, *Saccharomyces cerevisiae*; Uma, *Ustilago maydis*. Protein NCBI identification numbers are indicated.

RiMCO1, RiMCO3, RiMCO4, RiMCO5, RiMCO6 and RiMCO9 are predicted to have an N-terminal transmembrane domain, but lack the C-terminal transmembrane domain typical of ferroxidases. RiMCO8 was predicted to have three, one at the N-terminus and the other two at the C-terminus (Table S3 and Fig. S3).

These *in silico* analyses showed that although there is no homologue to Fet3-type ferroxidases sensu stricto in the *R. irregularis* genome, there are nevertheless several potential candidates that could have ferroxidase activity.

### RiMCOs *are differentially expressed in the intraradical and extraradical mycelium*

To further characterize the RiMCOs, we decided to compare *RiMCOs* gene expression in the IRM and ERM. Quantitative gene expression analysis was performed by real-time RT-PCR on ERM collected from the hyphal compartment of *R. irregularis* monoxenic cultures and on carrot mycorrhizal roots lacking ERM that were grown for two weeks in a densely colonized hyphal compartment of split Petri dishes. In the ERM, *RiMCO1, RiMOC4* and *RiMCO5* were the most highly expressed genes, followed by *RiMCO3. RiMCO6* expression was barely detected in the ERM. However, *RiMCO2* was the most highly expressed gene in the IRM. Transcript levels of *RiMCO1* and *RiMCO5* were approximately 10-fold higher in the ERM than in the IRM of the monoxenically grown carrot roots. No significant differences were observed between the expression levels of *RiMCO3, RiMCO4* and *RiMCO6* in both fungal structures *RiMCO2, RiMCO7, RiMCO8* and *RiMCO9* were not expressed or were below the detection limit in both fungal structures. (Fig. 5).

**Figure 5.**
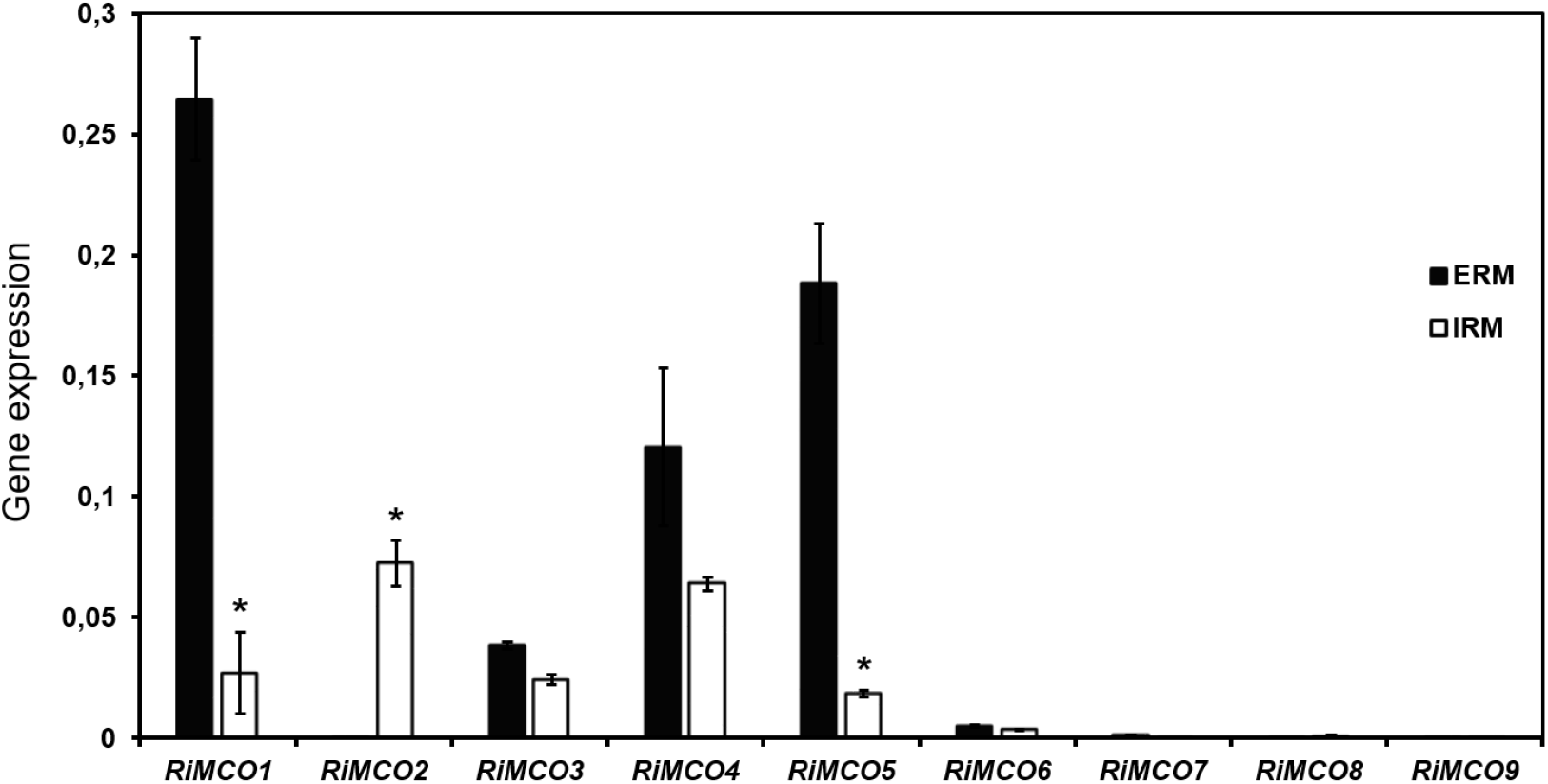
Relative expression of the *MCO* genes in extraradical mycelia (ERM) and intraradical (IRM) mycelia of *Rhizophagus irregularis*. *RiMCO* gene expression was assessed in ERM developed in monoxenic cultures (ERM) and *R. irregularis*-colonized carrot roots grown in monoxenic cultures and lacking ERM (IRM). Samples were normalized using the housekeeping gene *RiTEF*. Relative expression levels were calculated by the 2^-ΔCT^ method. Data are means +/-standard error. Asterisks show statistically significant differences (p<0.05) relative to the ERM, according to the Fisher’s LSD test.

### *Fe regulates* RiMCO1, RiMCO3 *and* RiMCO4 *gene expression*

Since fungal ferroxidases are transcriptionally regulated by Fe, in an attempt to identify the potential *R. irregularis* ferroxidases that interact with the Fe permeases RiFTR1 and RiFTR2, *RiMCOs* transcript levels were analyzed in ERM grown in the presence of different iron concentrations (Fig. 6). Relative to the ERM grown in control M media containing 0.045 mM Fe, *RiMCO1* and *RiMCO3* gene expression was observed to be 2.3- and 1.8-fold up-regulated, respectively, when the ERM was grown in medium lacking Fe (including addition of the Fe chelator ferrozine for 3 d). On the other hand, development of the fungus in the presence of 45 mM Fe induced a slight but statistically significant down-regulation of *RiMCO1* expression, and a 3.5-fold up-regulation of *RiMCO4* expression. Transcript levels of *RiMCO5* and *RiMCO6* were not significantly affected by the amount of Fe present in the culture medium. *RiMCO2, RiMCO7, RiMCO8* and *RiMCO9* were not expressed in our experimental conditions or were below the detection limit. *RiMCO1* and *RiMCO3* were the only *MCO* genes displaying a gene expression pattern typical of a high-affinity Fe transport system (Tamayo *et al*., 2018), suggesting that RiMCO1 and RiMCO3 might be partners of the Fe permeases RiFTR1 or RiFTR2. Moreover, *RiMCO4* gene was Fe-regulated, suggesting that it could also have a Fe-related role.

**Figure 6.**
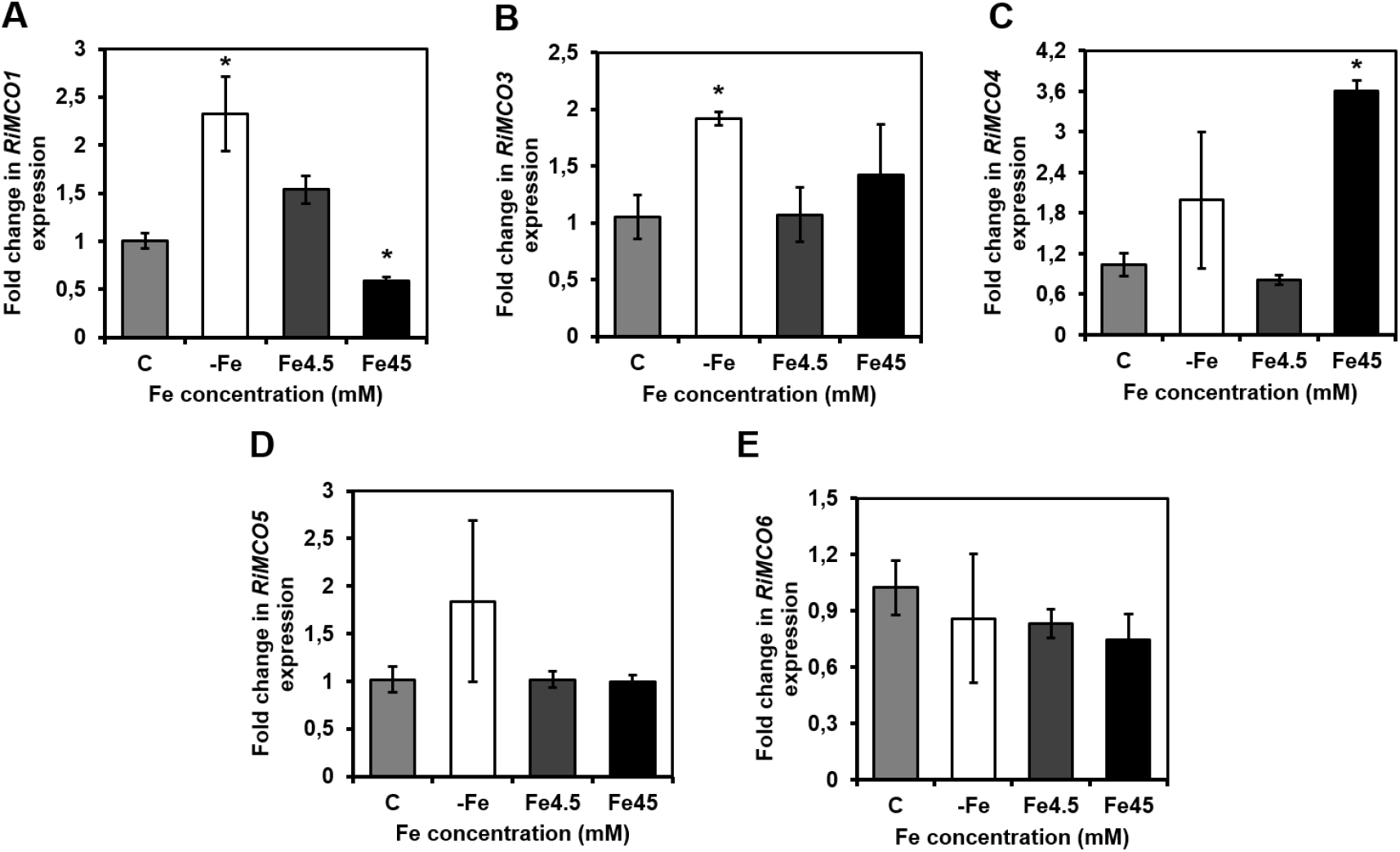
Effect of iron on the expression of the *Rhizophagus irregularis MCO* genes. *R. irregularis* was grown for 2 weeks in M-C media containing 0.045 mM Fe (control, C), 4.5 mM Fe (Fe4.5) or 45 mM Fe (Fe45), or in M-C without Fe and exposed for 3 days to ferrozine (-Fe). *RiMCO1* (A), *RiMCO3* (B), *RiMCO4* (C), *RiMCO5* (D) and *RiMCO6* (E) gene expression. Data were normalized using the housekeeping gene *RiTEF*. Relative expression levels were calculated by the 2-ΔΔCT method. Data are means +/-standard error. Asterisks show statistically significant differences (p<0.05) compared to the control value, according to the Fisher’s LSD test.

### *RiMCO1 and RiMCO3 enable Δ*fet3*yeast mutant to grow under Fe-limiting conditions*

To determine whether RiMCO1 and RiMCO3 could have a Fe-related function, we assessed if both proteins can restore the inability of a *S. cerevisiae* mutant lacking Fet3 ferroxidase activity (Δ*fet3*) to grow under Fe-limiting conditions. For this purpose, the *RiMCO1* and *RiMCO3* full length cDNAs were cloned into a yeast expression multicassette and the capability of the *RiMCO1*- and *RiMOC3*-expressing yeast mutant to grow under these conditions was determined. Relative to the untransformed Δ*fet3* cells, there was a better growth of the *RiMCO1*- and *RiMCO3*-expressing Δ*fet3* cells in Fe-limiting conditions (Fig. 7). Therefore, RiMCO1 and RiMCO3 enable the yeast mutant to take up iron.

**Figure 7.**
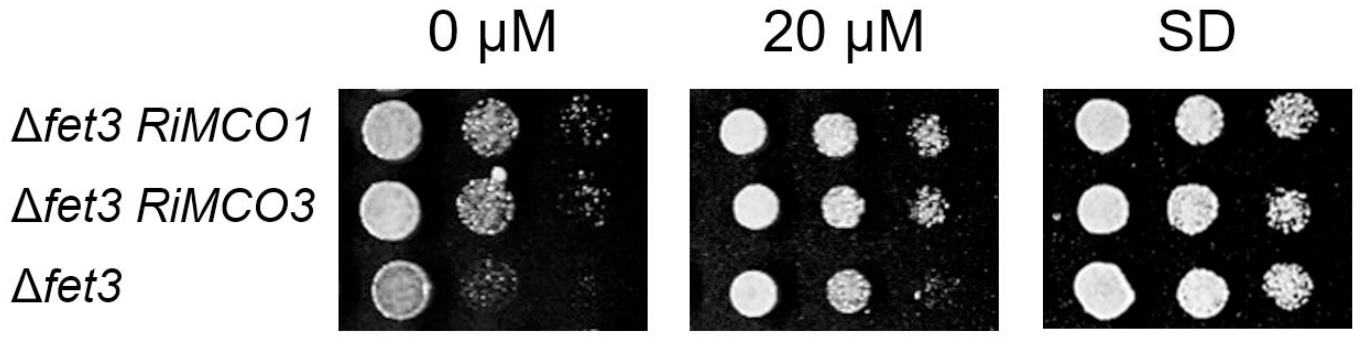
Analysis of the *in vivo* role of RiMCO1 and RiMCO3 in iron transport in yeast. Δ*fet3* cells untransformed or expressing *RiMCO1* or *RiMCO3* were examined for growth under Fe-limiting conditions (0 and 20 µM Fe). Control growth was observed in SD plates.

### RiFTR1 can take up Fe without the involvement of a ferroxidase partner in yeast

After the observation that RiMCO1 and RiMCO3 can contribute to Fe uptake in the heterologous system, we decided to examine which of them could act together with the plasma membrane Fe transporter RiFTR1 in the reductive Fe uptake pathway of *R. irregularis* (Tamayo *et al*., 2018), and with RiFTR2. To that end, the Δ*fet3*Δ*fet4* and Δ*fet3*Δ*fet4*Δ*ftr1* mutant strains, -lacking the ferroxidase Fet3 and the low-affinity Fe permease Fet4, and Fet3 and the low- and high-affinity Fe permeases Fet4 and Ftr1, respectively-, were transformed with plasmids containing *RiFTR1, RiFTR2, RiFTR1*-*R*i*MCO1, RiFTR1*-*RiMCO3, RiFTR2*-*RiMCO1* or *RiFTR2*-*RiMCO3*. Both mutant strains are unable to grow in Fe-limited media. Under Fe-limiting conditions, the lack of growth was restored in all the yeast cells transformed with any construct carrying the Fe transporter RiFTR1, but not with RiFTR2 (Fig. 8A and B).

**Figure 8.**
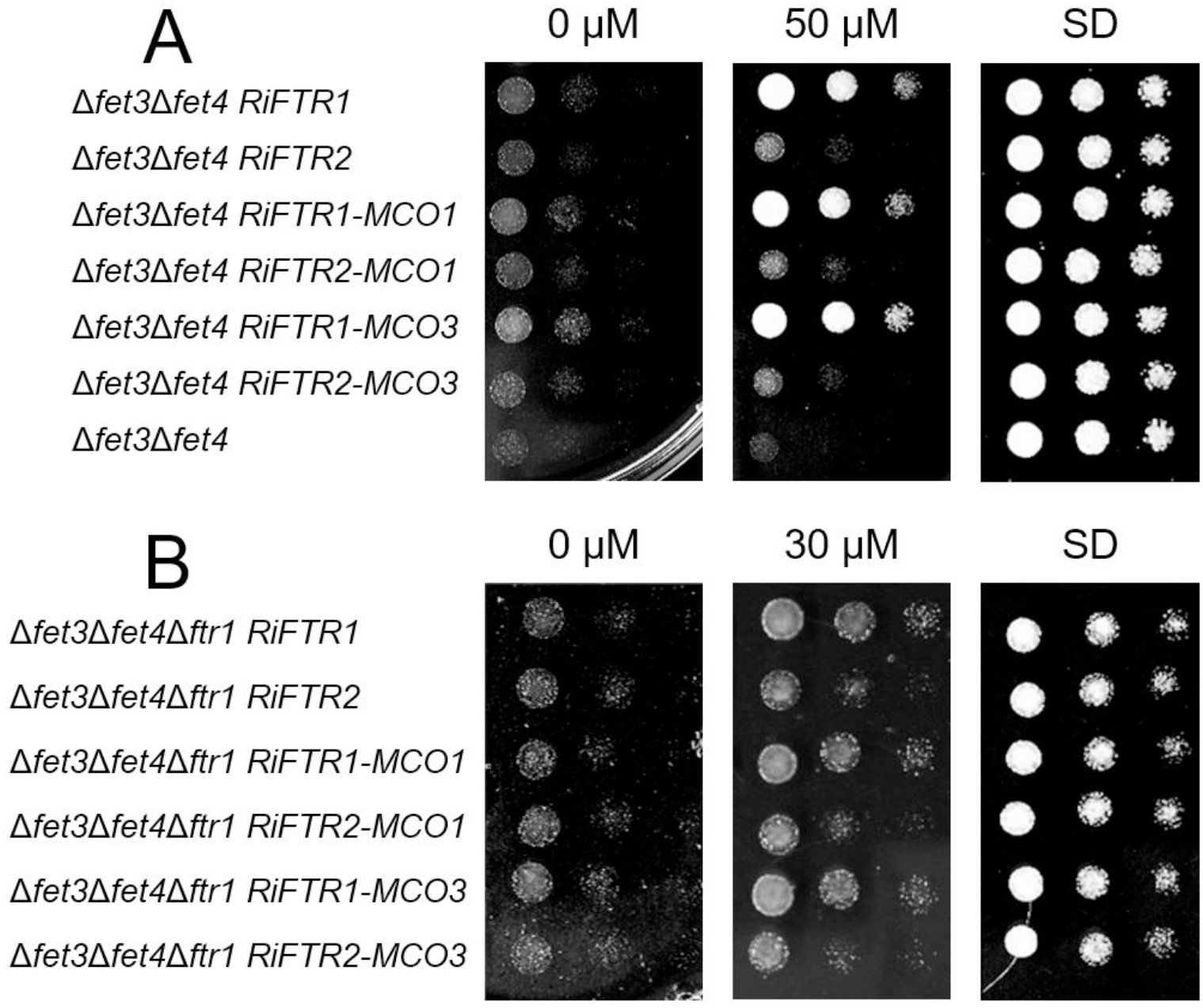
Analysis of the *in vivo* role of RiFTRs and RiMCOs in iron transport in yeast mutant strains lacking Fet3 and Fet4. Δ*fet3*Δ*fet4* **(A)** and Δ*fet3*Δ*fet4*Δ*ftr1* **(B)** yeast mutant strains were transformed with different constructs in order to search for the partner of the Fe transporter FTR1.

Co-expression of RiMCO1 or RiMCO3 did not help to improve the growth under Fe-limiting conditions. Furthermore, RiFTR1-mRuby2 fusion protein in the Δ*fet3*Δ*fet4* mutant strain was localized to the yeast plasma membrane (Fig. 9). These results indicate that this transporter is sufficient for the uptake of Fe under Fe-limiting conditions in yeast, without the need for the joint action of a ferroxidase partner.

**Figure 9.**
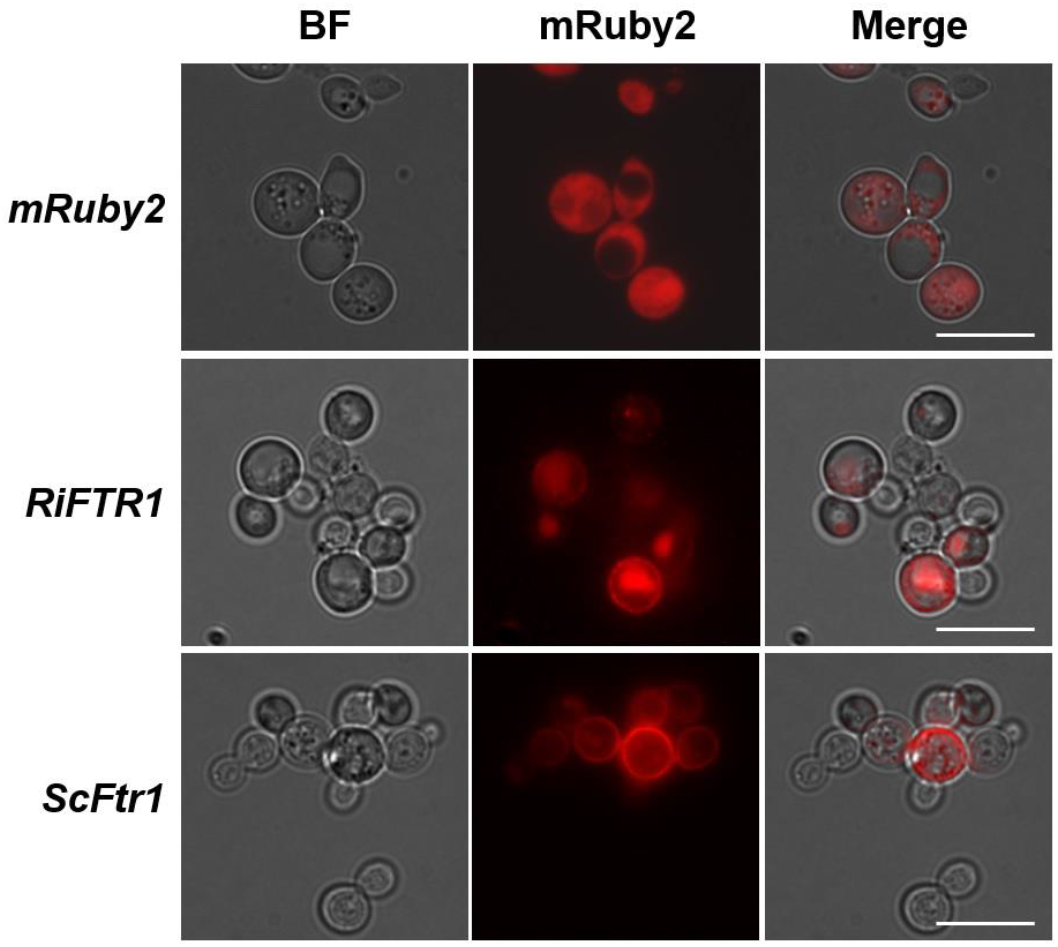
Analysis of the *in vivo* localization of RiFTR1 in a high-affinity reductive Fe pathway yeast mutant. Δ*fet3*Δ*fet4* cells transformed with a vector expressing *mRuby2* (first row), *RiFTR1-mRuby2* (second row) or *ScFtr1-mRuby2* (third row) were visualized with a Zeiss Axio fluorescent microscope. BF, bright field (first column); mRuby2, red channel (second column); Merge, combination of BF and red channels (third column).

## Discussion

Nine full-length sequences putatively encoding MCOs were identified in the genome of *R. irregularis*. However, in a search for potential ferroxidase partners for the Fe permeases RiFTR1 and RiFTR2, data obtained in this manuscript suggest that all RiMCOs belong to the ferroxidase/laccase branch of fungal MCOs and that *R. irregularis* therefore does not have any ferroxidase *sensu stricto*.

The number of putative *MCO* genes in *R. irregularis* is higher than those of *S. cerevisiae* (three MCOs; Hoegger *et al*., 2006), the basidiomycetes *P. chrysosporium* (five MCOs, Larrondo *et al*., 2004; Fernandez-Fueyo *et al*., 2012), *U. maydis* (six MCOs; Hoegger *et al*., 2006), *C. neoformans* (five MCOs; Kües and Rühl, 2011), and the mucoromycotina *R. oryzae* (five MCOs; Hoegger *et al*., 2006), but lower than those of the ascomycetes *Aspergillus niger* (sixteen MCOs, Tamayo-Ramos *et al*., 2011) and *Neurospora crassa* (eleven MCOs; Pöggeler, 2011). The number of MCOs in the AM fungi *R. irregularis, R. cerebriforme and R. clarus* is higher than in the closely phylogenetically related pathogen *R. oryzae*, the most common organism isolated from patients with mucoromycosis, but similar for instance to the number of the more phylogenetically distant ectomycorrhizal fungus *L. bicolor* (eleven MCOs; Courty *et al*., 2009). These data suggest that the relatively high number of *MCO* genes in AM fungi can be due to their lifestyle more than to their phylogenetic position. *In silico* protein analyses showed that the nine putative RiMCOs contain three cupredoxin-like domains typical of multicopper oxidases (Ajmad *et al*., 2022). The number of introns in the *MCO* sequences ranged between four and twelve and is lower than in basidiomycete *MCO* genes. For example, the *P. chrysosporium MCO* genes have 14-19 introns (Larrondo *et al*., 2004; Courty *et al*., 2009). Based on the phylogenetic tree and on intron distributions of the *R. irregularis* MCOs, the protein RiMCO6 seems to have an evolutionary origin different from the other MCOs. Intron positions of the *R. irregularis MCO* sequences defined five gene subfamilies, and it is remarkable that one gene of each subfamily is well expressed in the ERM.

To date, ferroxidase genes (coding Fet3-type ferroxidases or ferroxidases/laccases) have been found in all sequenced basidiomycetes except *Coprinopsis cinerea* (Kües and Rühl, 2011), whose genome also lacks *FTR1* homologs, but has a *sid1/sidA* gene involved in the biosynthesis of an iron-chelating siderophore involved in iron acquisition under conditions of low soil iron availability (Hoegger *et al*., 2006). Phylogenetic analyses of the MCO protein sequences of the reference fungi used in this study, including three basidiomycetes, three ascomycetes, one mucoromycotina species and three glomeromycotina fungi revealed that the RiMCO sequences of the three AM fungi of this study clustered together within the ferroxidases/laccases clade. Based on phylogenetic analyses of the sequenced genome to date, it seems that the lack of Fet3-ferroxidases *sensu stricto* might be true for all the glomeromycotina fungi. Therefore, we believe that *R. irregularis*, despite using a reductive pathway for iron uptake, is lacking a FET3-type ferroxidase ortholog in its genome, in contrast to the genomes of all the non-glomeromycotina fungi used in the phylogenetic study. Moreover, several RiMCOs are predicted to have an N-terminal transmembrane domain instead of the C-terminal transmembrane domain typical of ferroxidases. However, an N-terminal transmembrane domain was also identified in a novel *Acidomyces acidophilus* MCO oxidase that displays ferroxidase activity (Boonen *et al*., 2014). Moreover, the fact that the RiMCOs are lacking the key cysteine (C) in signature S2 shared in fungal laccases (Kumar *et al*., 2003) suggests that the *R. irregularis* MCOs do not exhibit laccase activity.

In several fungal species, at least one of the *Fet3* homologues is arranged in the genome with start-to-start codons in a mirrored tandem with the *Ftr1* homologue (Hoegger *et al*., 2006), which could indicate a common regulation and function of the genes in iron metabolism as proposed for the *Fet3/Ftr1* homologs of *S. pombe* (Askwith and Kaplan, 1997), *P. chrysosporium* (Larrondo *et al*., 2007) and *C. neoformans* (Jung *et al*., 2008) and shown to be present in many other fungi such as *A. fumigatus* (Schrettl *et al*., 2004), *F. graminearum* (Park *et al*., 2007), *U. maydis* (Eicchorn *et al*., 2006) and *L. bicolor* (Courty *et al*., 2009). We previously showed that the Fe permease RiFTR1 plays a key role in Fe acquisition (Tamayo *et al*., 2018). Such an arrangement in the *R. irregularis* genome does not exist, supporting the view that this fungus lacks a FET3-type ferroxidase. Nevertheless, we have found that *RiFTR1* is localized adjacent to a putative copper transporter (*CTR*) gene, suggesting that Fe metabolism could be connected to copper metabolism in this fungus, as demonstrated in other organisms (Bernal and Krämer, 2021; Crichton and Pierre, 2001).

The finding that *RiMCO1* and *RiMCO3* display Fe-regulated expression patterns typical of a high-affinity Fe transport system suggests that they might act as the RiFTR1 and RiFTR2 partners in *R. irregularis*. Heterologous expression of the selected candidates RiMCO1 and RiMCO3 in the yeast mutant lacking Fet3 ferroxidase activity (Δ*fet3*) partially restored growth under Fe-limiting conditions, indicating that both proteins might be able to interact with yeast Fe-permease Ftr1 and target to the plasma membrane, allowing the yeast mutant to take up iron. Therefore, these two proteins appear to be able to oxidize Fe in the heterologous system, allowing Fe(III) to be internalized by yeast Ftr1 subsequently.

Surprisingly, heterologous analyses in the triple yeast mutants lacking the ferroxidase fet3 and both the low- and high-affinity Fe uptake systems indicated that RiFTR1 is capable of transporting Fe under Fe-limiting conditions without the help of a ferroxidase partner. Co-expression of *RiMCO1* or *RiMCO3* together with either of the *R. irregularis* permeases RiFTR1 or RiFTR2 did not improve the growth of this triple mutant. These results suggest that they are not the partners of RiFTR1 in *R. irregularis*. Thus, RiFTR1 seems to be similar to the plant Fe-regulated transporter1 (IRT) system, capable of partially recovering the deficient growth of the Δ*fet3*Δ*fet4* yeast mutant under Fe-limited conditions (Eide *et al*., 1996; Quintana *et al*., 2022). Although this has not been found up to date in fungi, the fact that *R. irregularis* has no Fet3-ferroxidase encoded in its genome seems to support this possibility. These data suggest that the reductive Fe uptake pathway evolved by AM fungi is more similar to the reduction strategy of plants than to that of fungi. To acquire Fe from soil, higher plants have evolved two different strategies: a reduction strategy (Strategy I) employed by dicots and non-graminaceous monocots and a chelation strategy (Strategy II) used by graminaceous species (Marschner and Römheld, 1994). In the reduction strategy, once that Fe is solubilized by the action of a H+-ATPase, Fe(III) is reduced to the more soluble form Fe(II), which is subsequently transported into the root epidermal cells via high-affinity iron transporters from the Zrt/Irt-like protein (ZIP) family. Therefore, our data suggest that Fe transport across the *R. irregularis* ERM plasma membrane through the Fe permease RiFTR1 occurs in the form of Fe(II) and that does not require the concerted action of a ferroxidase, as it has been described in fungi that employ a reductive Fe uptake pathway (Philpott, 2006). Given that AM fungi facilitated the initial colonization of land by plants by providing them inorganic nutrients and moisture from the substrate in exchange for carbon compounds (Selosse *et al*., 2015), that roots evolved after the transition to lands (Bundrett, 2002) and the more than 400 million years of coevolution of AM fungi and plants (Strullu-Derrien *et al*., 2014), it is not striking that the AM fungi and plants use the same Fe uptake strategy to take up Fe from the soil solution. In fact, it could be hypothesized plants exploited the reductive Fe uptake strategy previously evolved by AM fungi.

The biological functions of MCOs in nature are diverse and they have been involved in a wide variety of processes, such as pigmentation, stress response to environmental challenges, fungal morphogenesis and plant-fungal interactions (Kües and Rühl, 2011). Given that the ERM exposed to 45 mM Fe presented a brownish colour (data not shown), up-regulation of *RiMCO4* under these conditions suggests that its encoded protein might be involved in fungal pigmentation and senescence, as it has been suggested for some MCOs of the white rot fungus *Lentinula edodes* (Sakamoto *et al*., 2015). Differential expression of the *RiSMF* genes in the IRM and ERM agrees with previous observations in the pathogenic fungus of maize *Setosphaeria turcica*. Expression of the nine MCOs of *S. turcica* were differentially expressed in the different developmental stages of the fungus and it was hypothesized that in this fungus MCOs play different roles during fungal growth and infection processes (Liu *et al*., 2019). As it has been reported in *L. edodes*, the *R. irregularis* MCOs might be involved in hyphal morphogenesis and cell wall synthesis (Nakade *et al*., 2011).The higher transcript level of *RiMCO1* and *RiMCO5* in the ERM than in the IRM indicates that they could have a main role in the extraradical mycelium of the fungus. *RiMCO2*, whose expression was under the limit of detection in the ERM, was highly expressed in the IRM, suggesting a role in the functioning of fungal symbiotic tissues.

In conclusion, this study shows that the AM fungus *R. irregularis* has at least nine MCO members in its genome. Moreover, we showed that the Fe permease RiFTR1 is able to take up Fe under Fe-limiting conditions without the need for a ferroxidase partner. Further studies are required to ascertain the biological function of the *R. irregularis* MCOs.

## Acknowledgements

This research was funded by Spanish MCIN/AEI/10.13039/501100011033/ and FEDER Una manera de hacer Europa, grant numbers AGL2015-67098-R and RTI2018-098756-B-I00 and TUM funding. Elisabeth Tamayo was supported by a PhD contract (JAEPre054) from CSIC. We are grateful to Drs. A. Dancis and S. Knight (University of Pennsylvania) for donation of 42B, YPHfa and DDY4 yeast mutant strains and the vector containing *S. cerevisiae Ftr1* and to Dr. D. Eide (University of Wisconsin-Madison) for donation of DEY1530 yeast mutant strain and the vector containing *S. cerevisiae Fet3*. We also thank M. Rüllke, M.Sc. (Technical University of Munich) for his help with the design of the Golden Gate cloning strategy.

